# CircGLCE alleviates intervertebral disc degeneration by regulating apoptosis and matrix degradation through the targeting of miR-587/STAP1

**DOI:** 10.1101/2020.03.06.981621

**Authors:** Zhonghui Chen, Weibing Zhang, Ming Deng, Yan Zhou, Yaming Li

## Abstract

**Background:** Intervertebral disc degeneration (IDD) can induce profound global socioeconomic burdens. Recent studies have suggested that circular RNAs might have crucial functions in the progression of IDD. The purpose of this study was to identify a specific circular RNA and to investigate its regulatory mechanism in IDD.

**Methods:** CircGLCE was selected after microarray analyses and was further analysed by RT-qPCR and FISH. After silencing CircGLCE, its function was assessed with RT-qPCR, immunofluorescence analysis and flow cytometry. Based on Sanger sequencing, miR-587 was identified as a direct target of CircGLCE, and it was further examined with RNA pulldown assays, RT-qPCR, dual luciferase assays and FISH. After silencing CircGLCE or miR-587, western blotting, immunofluorescence analysis, and flow cytometry were conducted. STAP1 was assessed by RT-qPCR and luciferase assay, and experiments with silenced and overexpressed miR-587 were performed. A rescue experiment was also included. In an IDD rat model, the in vivo effects of overexpressing CircGLCE on IDD were analysed with imaging techniques, TUNEL staining, FISH, western blotting, H&E staining and immunohistochemistry.

**Results:** CircGLCE was found to stably exist in the cytoplasm of nucleus pulposus (NP) cells. It was downregulated in IDD. Knockdown of CircGLCE promoted apoptosis and induced the expression of matrix-degrading enzymes in NP cells. CircGLCE served as a miR-587 sponge in NP cells. Inhibiting miR-587 counteracted the IDD-enhancing effect caused by silencing CircGLCE. STAP1 served as the miRNA target that mediated the functions of miR-587. Overexpressing CircGLCE alleviated IDD in vivo.

**Conclusions:** CircGLCE attenuates IDD by regulating the apoptosis of NP cells and by regulating ECM degradation through the targeting of miR-587/STAP1. CircGLCE may be a potential therapeutic target for IDD treatments.

## 1. Introduction

As the leading illness threatening public health and well-being,^1^ various degrees of lower back pain is known to occur in approximately 80% of the global population, and it is the direct cause of chronic disability among 10% of sufferers.^2^ The major underlying physiological process involved in lower back pain is intervertebral disc degeneration (IDD), which leads to progressive structural abnormality of the lumbar spine and accelerated ageing of the intervertebral discs.^3, 4^ Anatomically, the gelatinous nucleus pulposus (NP) of the intervertebral disc is surrounded by annulus fibrosis (AF).^5^ NP cells produce most of the major components of the extracellular matrix, including Collagen II and aggrecan, which are essential for the integrity of disc structure.^6, 7^ Therefore, once NP cells are dysfunctional due to the abnormal cellular activities caused by IDD, such as the increased release of proinflammatory factors and apoptosis, the degradation of the ECM is induced, which further facilitates IDD.^8-10^ Accordingly, the inhibition of these abnormal activities by NP cells is the key to preventing the progression of IDD.^11, 12^

Numerous biological and genetic regulators have been associated with IDD pathogenesis^13-16^; nonetheless, recently discovered circular RNAs (circRNAs) have also been suggested to be crucial elements based on accumulating evidence.^17-19^ The single-stranded non-coding RNA sequences are endogenously expressed from the genome, and they form closed loops through covalent bonding. Basically, circRNAs serve as competing endogenous mRNAs (ceRNAs) by directly binding to miRNAs to counteract the posttranscriptional repression of target mRNAs mediated by miRNAs; in this process, the ceRNA is referred to as a miRNA sponge.^20^ Subsequently, the downstream signalling pathways associated with miRNAs are prone to regulation by specific circRNAs. It has been reported that circRNAs are involved in multiple pivotal biological processes, including cell proliferation, cell differentiation, apoptosis, and metabolism,^17-19^ and circRNA expression patterns are associated with a variety of diseases, such as malignancies,^21, 22^ immune disorders,^23^ cardiovascular conditions,^24^ and neurological degeneration.^25^ To explore the role of circRNAs in IDD, a differential expression profile was produced and was analysed with a bioinformatic approach; the data revealed a promising candidate, CircGLCE.^26^ In vivo experiments were performed using a rat tail model, which provides readily detected intervertebral disc degeneration through mechanical injury.^27^ CircGLCE knockout and mutant rat models enabled the investigation of the effects of eliminating CircGLCE. Probing of downstream targets was performed to provide further clarity on the pathogenetic mechanism of IDD and to identify novel therapeutic targets.

## 2. Materials and Methods

### 2.1 Patient samples

A total of 194 patients (58.6 ±6.1 years, 75 men and 119 women) were retrospectively recruited for degenerative disc sample collection by discectomy, which was a part of the surgical treatment procedures. The application of surgical treatment depended on the following conditions: (1) the failure of conservative treatments and (2) the diagnosis of progressive neurologic disorders, including cauda equina syndrome and progressive muscle weakness. The exclusion criteria were predominantly concerned with predispositions for vertebral structural abnormalities, inflammatory conditions or other neuropathological events, such as ankylosing spondylitis, diffuse idiopathic skeletal hyperostosis, isthmic or degenerative spondylolisthesis and lumbar stenosis. A total of 152 patients were recruited to be used as controls, which involved collecting NP tissue samples taken from fresh traumatic lumbar fracture as the result of anterior decompressive surgery. The experimental group and the control group were matched for age distribution and sex. For all of the participants, the lumbar spine was scanned by magnetic resonance imaging, which was a part of routine clinical procedures. Based on the T2-weighted images, the severity of IDD was evaluated with the Pfirrmann grading system. The distribution of the degree of degeneration in IDD samples was as follows: grade 2 in 15 samples, grade 3 in 44, grade 4 in 85, and grade 5 in 50. Seventy-two of the 194 samples were obtained from the level of L4/L5, and 122 were from L5/S1. This study protocol conformed to the globally accepted regulations on clinical studies involving human data, and approval was conferred by the ethics committee of Renmin Hospital of Wuhan University. Written informed consent was obtained from all of the participants before using the samples. The experiments and reporting steps were performed in exact accordance with the designed protocol and the ethical requirements.

### 2.2 Surgically induced IDD animal model

The IDD model was established in WT C57BL/6 mice (12 weeks old) by AF needle puncture. In brief, the general anaesthesia used was ketamine (100 mg/kg). A sagittal small skin incision was performed from Co6 to Co8 to help locate the disc position for needle insertion in the tail. Subsequently, Co6–Co7 coccygeal discs were punctured using a needle. The syringe needle was inserted into the Co6–Co7 disc along the vertical direction and then rotated in the axial direction by 180° and then was held for 10 s. The puncture was made parallel to the endplates through the AF into the NP using a 31-G needle, which was inserted 1.5 mm into the disc to depressurize the nucleus. The other segments were left undisturbed for contrast. The wound was closed. For the therapeutic experiment, discs were harvested at 12 weeks post surgery from WT mice. All procedures and protocols were approved by Renmin Hospital of Wuhan University.

### 2.3 Primary culture of nucleus pulposus (NP) cells

Before culturing human NP cells, the specimen taken from the intervertebral discs required purification, which involved 3 rinses with phosphate-buffered saline (PBS; Gibco, NY), division of the tissue into tiny pieces, and digestion with a solution containing type II collagenase (0.2% w/v; Gibco) and trypsin (0.25% w/v; Gibco); then the mix was subjected to a 3-hour incubation in PBS at 37 °C with constant shaking before being filtered. The mesh size of the filters was 70 μm (BD, NJ). Dulbecco’s modified Eagle’s medium was used for primary culture, and it was supplemented with Ham’s F-12 Nutrient Mixture (DMEM-F12; Gibco), 50 μg/mL streptomycin (Gibco), 50 U/mL penicillin, and 20% (v/v) foetal bovine serum (FBS; Gibco). The medium mixture was poured into 10 cm culture dishes and then was incubated in the presence of 5% CO_2_. Once approximately 80% confluence was achieved following trypsin treatment, the acquired NP cells were subcultured into 6 cm culture dishes at a density of 2.5×10^5^ cells per dish. Passages 1 and 2 were considered suitable for further testing in this study.

### 2.4 RNA extraction from tissue samples

At the time of isolation from the body, the AF and NP tissue were carefully separated from each other and then were chopped into small pieces (1-3 mm^3^) under the microscope using a scalpel. Then, samples were snap frozen in liquid nitrogen within 40 min of removal from the patient and stored at −80°C until RNA extraction. RNA extraction was performed within one week of sample collection. Frozen tissue biopsies were ground into a powder using a mortar and pestle and then were homogenized in Qiazol reagent (Qiagen, Hilden, Germany). Total RNA was extracted using a miRNeasy Mini kit (Qiagen, Venlo, The Netherlands) and was quantified using a NanoDrop 2000 spectrophotometer (Thermo Fischer Scientific, Wilmington, Delaware). RNA integrity was assessed using an Agilent Bioanalyzer DNA 1000 chip (Invitrogen, Carlsbad, CA). The RIN values were no less than 9.6.

### 2.5 Microarray analysis

Total RNA was extracted with TRIzol reagent (Invitrogen, MA), and then the RNA was purified with an RNeasy Mini kit (Qiagen, Germany). Six pairs of degenerative NP samples (100 mg each) and matched controls were tested. Each RNA sample was processed with fluorescence-labelling to prepare complimentary RNA (cRNA) targets that could be detected by an SBC human ceRNA array V1.0 (4□×□180K, Agilent Technologies). The microarray kit contained 88,371 circRNA probes and 18,853 mRNA probes. The processed cRNA transcripts were transferred onto slides for hybridization. Subsequently, the slides were read by an Agilent Microarray Scanner (Agilent, CA). Feature Extraction software 10.7 (Agilent Technologies) was utilized for data acquisition. The normalization of raw data relied on the quantile algorithm of the R program within the LIMMA package. The standards used to identify expressed transcripts with differential expression levels between the IDD samples and controls were set up to be fold change ≥2 or ≤−2, based on p-value□≤□0.05.

### 2.6 RNA isolation and RT-qPCR

Total RNA was extracted with TRIzol Reagent (Ambion, CA). The concentration and purity of the extracted RNA was measured by a NanoDrop Spectrophotometer (Thermo, MA) and an Agilent 2100 Bioanalyzer (Agilent, CA). To quantify specific miRNA and circRNA content, a TaqMan MiRNA assay kit (Applied Biosystems, CA), an iScript Select cDNA Synthesis kit (Bio-Rad, CA), and an iQSupermix kit (Bio-Rad, CA) were used to enable target-specific detection, and the procedures complied with the manufacturer’s instructions. To quantify mRNA levels, an iScript Select cDNA Synthesis kit was also adopted to produce complimentary DNA transcripts with oligo-dT primers. Based on the primer set specialized for target mRNA, iQSYBR Green mix (Bio-Rad) was applied to enable fragment amplification by real-time quantitative reverse transcription PCR (RT-qPCR). The reactions were carried out on a PCR flatform (Applied Biosystems) that also enabled capturing of measurements. The raw quantitative results were normalized to the level of β-Actin, which was followed by data processing with the comparative Ct (ΔΔCt) method (2 ^−ΔΔCt^ with logarithm transformation) to estimate the relative expression levels.^28^

### 2.7 Small RNA library

Total RNA extraction was performed using TRIzol reagent (Invitrogen, MA). Six pairs of degenerative NP samples (100 mg each) and matched controls were tested. An Agilent 2100 Bioanalyzer (Agilent, CA) was utilized to examine RNA integrity (RIN). The library construction was performed using an Illumina TruSeq small RNA Sample Prep kit (Illumina, CA) and 1 μg of total RNA following the manufacturer’s instructions. Fifteen cycles of PCR were carried out. The quality of the RNA product was examined with a High Sensitivity DNA Chip and an Agilent 2100 Bioanalyzer (Agilent Technologies, CA), and the concentration was assessed by qPCR using a KAPA Library Quantification kit (KAPA Biosystems, CA). Each library was adjusted to a concentration of 20 pM and was sequenced with a MiSeq Reagent kit v3 for 150 cycles by an Illumina MiSeq Sequencing System (Illumina). An efficiency of 25 million sequence reads per flow cell was achieved.

### 2.8 Bioinformatics analysis

The miRNA targets of CircGLCE were predicted by two bioinformatic programs: TargetScan (http://www.targetscan.org/vert_71/) and miRanda (http://www.microrna.org/microrna/home.do, total score > 145). GO and KEGG analyses were performed to predict the influences of the downregulated mRNAs in the HCs using the DAVID bioinformatics program (https://david.ncifcrf.gov/tools.jsp). The potential target genes of miR-587 were predicted by TargetScan (http://www.targetscan.org/vert_71/), miRDB (http://mirdb.org) and RNA-seq results. The potential Ago2 and other RNA binding protein binding sites of CircGLCE were obtained using the circRNA online tool (https://circinteractome.nia.nih.gov/bin/circsearchTest).

### 2.9 Pull-down assay with biotinylated CircGLCE probe

A total of 107 NP cells were washed with PBS and then were cross-linked by ultraviolet irradiation at 254 nm. Cells were lysed with 1 ml of lysis buffer and then were fully homogenized with a 0.4 mm syringe. One biotinylated antisense probe (0.2 nmol) targeting the adaptor sequence or Lac Z probes (Control probes) was added to the CircGLCE-Pull down system. The probes were denatured at 65 °C for 10 min and hybridized at room temperature for 2 h before adding 200 µl of streptavidin-coated magnetic beads. Nonspecifically bound RNAs were removed by washing, and TRIzol reagent was used to recover miRNAs specifically interacting with CircGLCE. PCR and RT-qPCR were used to analyse binding strength after reverse transcribing the miRNAs.

### 2.10 Western blot analysis

The cells were lysed using RIPA buffer (Sigma, St. Louis, MO, USA). Total protein was separated by 12% sodium dodecyl sulphate-polyacrylamide (SDS-PAGE) electrophoresis and then was transferred to nitrocellulose membranes (Millipore, Billerica, MA, USA), which was followed by incubation with antibodies against STAP1, Aggrecan, Col II, ADAMTS5, MMP13 and β-actin (Cell Signalling Technology, Boston, USA) overnight at 4 °C. After being rinsed three times, the membranes were further incubated with an HRP-conjugated anti-IgG for 1 h at 37°C. The protein was detected using an ECL system (Amersham Pharmacia, Piscataway, NJ, USA) and was analysed using Quantity One software (BIO-RAD, USA).

### 2.11 Northern blot analysis

Northern blot analysis was performed with a northern blot kit (Ambion, USA). Briefly, total RNA (∼30 μg) from NP cells was denatured in formaldehyde and then was electrophoresed in a 1% agarose-formaldehyde gel. The RNAs were then transferred onto a Hybond-N + nylon membrane (Beyotime, China) and were hybridized with biotin-labelled DNA probes. A biotin chromogenic detection kit (Thermo Scientific, USA) was used to detect the bound RNAs. Finally, the membranes were exposed and analysed using Image Lab software (Bio Rad, USA).

### 2.12 RNA fluorescent in situ hybridization (FISH)

FISH assays were performed in NP cells or NP tissues.^29^ Cy3-labelled CircGLCE probes and 488-labelled locked nucleic acid miR-587 probes were designed and synthesized by RiboBio (Guangzhou, China). The signals of the probes were detected by a Fluorescent in situ hybridization kit (RiboBio, Guangzhou, China) according to the manufacturer’s instructions. The images were acquired on a Nikon A1Si laser scanning confocal microscope (Nikon Instruments Inc., Japan). For in vivo FISH, tissue sections were deparaffinized, rehydrated, and permeabilized by 0.8% pepsin treatment at 37°C for 30 min before hybridization.

### 2.13 Transfection reporter assay

NP cells were seeded in 96-well plates and were cultured to 70%-80% confluence before transfection. For CircGLCE and miR-587 vector generation, either wild-type or mutant CircGLCE fragments (489 bp) were inserted into the Xba1 restriction sites of the pGL3-firefly luciferase vector. A total of 600 ng of plasmid (CircGLCE-wt and CircGLCE-mut) and 20 nmol miR-587 and NC were transfected. For STAP1 and miR-587, either wild-type or mutant STAP1 3’UTR fragments (116 bp) were inserted into the Xba1 restriction sites of the pGL3-firefly luciferase vector (Genechem, Shanghai, China). Five hundred nanograms of STAP1 3’UTR-wt and STAP1 3’UTR-mut plasmids and 20 nmol miR-587 and NC plasmids were transfected. After 48 h of incubation, a Promega Dual-Luciferase system was used to detect firefly and Renilla luciferase activities. Using 100 ml of Luciferase Assay Reagent II (LAR II) (Luciferase Assay Reagent, Promega) and, subsequently, 20 ml of lysis buffer, firefly luciferase activities were measured to provide an internal reference, and then Renilla luciferase activities were measured after the addition of 100 ml of Stop & Glo® Reagent (Luciferase Assay Reagent, Promega). Finally, the differences between firefly and Renilla luciferase activities were calculated to determine relative luciferase activity.

### 2.14 Flow cytometry analysis

The cells were stained with 5 µL of Annexin V-FITC and 10 µL of PI. Cells that were positively stained with Annexin V-FITC and negatively stained for PI were considered apoptotic. Cells that were positively stained for both Annexin V-FITC and PI were considered necrotic. Flow cytometry analysis was conducted with EpicsAltra (Beckman Coulter, CA, USA) to isolate the cells and determine their proportions.

### 2.15 CircGLCE knockdown with an adenoviral vector

For construction of recombinant adenoviral shCircGLCE, oligonucleotides with the CircGLCE targeting sequences were used for the cloning of small hairpin RNA (shRNA)-encoding sequences into the pDC311-U6-MCMV-EGFP vector (purchased from Hanbio Co. Ltd, Shanghai, China). For adenovirus miR-587 sponge construction, self-complementary DNA oligonucleotides encompassing the sequence of miRNA miR-587 were chemically synthesized, including overhang sequences from a 5’BamH1- and a 3’EcoRI-restriction site. Annealed oligonucleotides were directionally cloned into the BamH1/EcoRI-digested pDC311-U6-MCMV-EGFP vector (purchased from Hanbio Co. Ltd, Shanghai, China). pDC311-shCircGLCE or pDC311-mir-587-sponge and pBHGlox E1,3Cre were cotransfected into HEK-293 cells using LipoFiterTM transfection reagent (Hanbio, Shanghai, China) to generate recombinant adenoviruses (Ad-shCircGLCE). Adenoviruses harbouring green fluorescent protein (Ad-GFP) were used as a control. Ad-shCircGLCE or pDC311-mir587-sponge and Ad-GFP were propagated in HEK-293 cells. The propagated recombinant adenoviruses in the HEK-293 cells were purified, and the virus titer was measured by plaque assays. The stock solutions of Ad-shCircGLCE or pDC311-mir587-sponge and Ad-GFP were maintained at 1×10^10^ plaque formation units (PFUs)/ml.

### 2.16 Immunohistochemistry

Tissue sections taken from the intervertebral discs of mice were deparaffinized and boiled in 10 mM citrate buffer (pH 6.0) for antigen retrieval. Endogenous peroxidase was blocked by 3% H_2_O_2_. Then, slides were blocked in serum, incubated with anti-MMP13 (diluted 1:500, Abcam Cat# ab39012, RRID: AB_776416) at 4 °C overnight, incubated with an anti-rabbit secondary antibody, and visualized with diaminobenzidine (Sigma). A negative control experiment was also performed. The images of IHC staining were captured with a microscope (Olympus).

### 2.17 Immunofluorescence staining

Cultured human NP cells were rinsed with PBS three times, fixed with 4% formaldehyde for 15 min, and permeabilized with 0.3% Triton-X100 (Sigma, CO, USA) for 10 min. Subsequently, the cells were blocked with 6% BSA for 1 h and then were incubated overnight at 4°C with primary antibodies, such as anti-MMP13 (diluted 1:200, Abcam Cat# ab39012, RRID: AB_776416), anti-COL II (diluted 1:1000, Abcam Cat# ab34712, RRID: AB_731688), anti-Aggrecan (diluted 1:200, Abcam Cat# ab36861, RRID: AB_722655) or anti-ADAMTS-5 (diluted 1:500, Abcam Cat# ab41037, RRID: AB_2222327). Then, the cells were incubated with Alexa Flour 488-labelled goat anti-mouse IgG (Invitrogen 1:200; Invitrogen, OR, USA) for 1 h at room temperature. Cells were analysed using an Olympus DP72 fluorescence microscope (Olympus, Tokyo, Japan). To detect NP cell apoptosis, TUNEL staining was performed using a kit based on the manufacturer’s instructions (Promega, Fitchburg, WI, USA).

### 2.18 Injection of CircGLCE

AAV CircGLCE wt and mut constructs were generated and packaged by Vigene Biosciences. One week after the initial surgery, a total of 18 IDD male mice were randomly divided into three groups (control injection, CircGLCE injection, and CircGLCE-mut injection; n=6 per group). The solution containing experimental or control virus (approximately 1*10E13 vg/ml) overexpressing human CircGLCE was slowly injected into discs. Rats were treated with the local delivery of control, CircGLCE, or CircGLCE-mut on days 1, 7, and 14 after IDD surgery. At 12 weeks after surgery, discs were harvested for histological and radiographic evaluation in each group.

### 2.19 Radiographic evaluation

Radiographs were taken at 6 and 12 weeks after the injection. After the last radiography, MRI was performed on all rats using a 7.0 T animal-specific MRI system (Bruker Pharmascan, Ettlingen, Germany). T2-weighted sections in the median sagittal plane were obtained using the following settings: a fast spin echo (SE) sequence with a time to repetition (TR) of 3,000 ms and a time to echo (TE) of 70 ms; the slice thickness was 0.5 mm, and the gap was 0 mm. Pfirrmann classification was used to assess the degree of IDD degeneration. The average score of the punctured IDD was calculated as the amount of degeneration for each rat. The change in IVD height was evaluated by the disc height index (DHI). Measurements of internal control discs were carried out together with their corresponding punctured discs. Disc height and the adjacent vertebral body heights were measured from the midline as 25% of the disc’s width from the midline on either side. The DHI was expressed as the mean of the 3 measurements from midline to the boundary of the central 50% of disc width divided by the mean of the 2 adjacent vertebral body heights. Changes in the DHI of punctured discs were expressed as a percentage (%DHI=post-punctured DHI/pre-punctured DHI x100).

### 2.20 Histological evaluation

The discs from mice were fixed in 10% neutral-buffered formalin for 1 week, decalcified in EDTA for 2 weeks, paraffin-embedded, and carefully cut to generate 5-μm thick sections. Midsagittal sections were stained with haematoxylin and eosin. The histological images were analysed using an Olympus BX51 microscope (Olympus Centre Valley, PA, USA). Based on a literature review of disc degeneration studies,^30-35^ a modified histologic grading system was developed. More specifically, the cellularity and morphology of the AF, NP, and the border between the two structures were examined.^36^ The scale is based on 5 categories of degenerative changes with scores ranging from 0 points (0 in each category) for a normal disc to 15 points (3 in each category) for a severely degenerated disc.

### 2.21 Statistical analysis

Each test was repeated in triplicate, and the results were estimated as the mean±SEM. All of the calculation steps performed in GraphPad Prism 8 (GraphPad Software, CA). Unpaired two-tailed Student’s t tests were applied to evaluate the difference between two sets of data. Among multiple groups, one-way analysis of variance (ANOVA) coupled with Tukey’s post hoc test was applied to evaluate the difference. The p value accepted as a significant difference was set as p<0.05.

## 3. Results

### 3.1 CircGLCE stably exists in the cytoplasm of NP cells but is downregulated in IDD

In the heat map generated by microarray analysis on 6 IDD samples and the paired controls, as shown in Fig. 1a, circPRIM2, circCCT4, circLRP5, circGLCE and circDHX8 were observed to be significantly dysregulated. Then, further validation was performed by RT-qPCR using another cohort that included 31 IDD samples and 26 normal controls, which were randomly selected from 194 degenerative NP samples and 152 controls. Finally, as shown in Fig. 1b, only circGLCE was found to be significantly downregulated in IDD patients compared with its levels in controls (P<0.001). No significant difference was observed between IDD and controls with respect to circPRIM2, circCCT4, circLRP5 and circDHX8 (P=0.34, 0.18, 0.09 and 0.29, respectively; Table 1). Thus, we focused on circGLCE for further study. The FISH results also confirmed the differential expression levels of CircGLCE when comparing an IDD sample with a control sample (Fig. 1c). As shown in Fig. 1d, the head-to-tail splicing of CircGLCE that was determined by Sanger sequencing applied to the analysis of its RT-PCR transcripts. To exclude the interference of back-splicing,^37^ convergent primers were applied to amplify GLCE mRNA, while divergent primers were introduced to amplify CircGLCE. As a result of amplification with divergent primers, the band representing CircGLCE was present in the cDNA but absent in the genomic DNA (gDNA), as shown in Fig. 1e. With the CircGLCE-specific probe targeting the junction, the bands of 842 nt were also shown in northern blotting experiments, as expected (Fig. 1f and 1g), regardless of RNase R treatment. Therefore, the splicing of CircGLCE was confirmed based on the RNase R resistance of the transcripts, which was in contrast to the decreased GLCE mRNA level caused by RNase R. In addition, the results of FISH (Fig. 1h) and RT-qPCR (Fig. 1i) experiments confirmed that the cytoplasmic CircGLCE was detected at a remarkable level in NP cells.

**Table 1.** Differentially expressed circRNAs in NP cells in IDD patients compared with the controls in both the training set and the validation set

| miRNAs | Training set |  | Validation set |  |
| --- | --- | --- | --- | --- |
|  | Fold change | P Value | Fold change | P Value |
| **Up-regulated** |  |  |  |  |
| Hsa-circPL9 | 5.8 | 0.16 | - | - |
| **Hsa-circPRIM2** | 4.9 | **0.006\*\*** | 4.7 | 0.34 |
| Hsa-circTPP2 | 6.8 | 0.21 | - | - |
| Hsa-circMYO1E | 5.2 | 0.35 | - | - |
| Hsa-circMAP7 | 8.9 | 0.06 | - | - |
| Hsa-circSLC20A2 | 10.8 | 0.07 | - | - |
| **Hsa-circCCT4** | 13.9 | **0.005\*\*** | 12.6 | 0.18 |
| Hsa-circEXOC6 | 7.2 | 0.33 | - | - |
| Hsa-circFOXN2 | 6.3 | 0.05 | - | - |
| Hsa-circSLC35D2 | 4.2 | 0.09 | - | - |
| Hsa-circITGB5 | 6.1 | 0.21 | - | - |
| Hsa-circCOL17A1 | 4.6 | 0.07 | - | - |
| Hsa-circSF3B1 | 3.5 | 0.17 | - | - |
| **Hsa-circLRP5** | 7.6 | **0.001\*\*** | 7.3 | 0.09 |
| Hsa-circSDAD1 | 3.9 | 0.26 | - | - |
| Hsa-circTMCO4 | 4.2 | 0.44 | - | - |
| Hsa-circLAMB1 | 8.6 | 0.51 | - | - |
| Hsa-circPIK3C3 | 4.2 | 0.1 | - | - |
| Hsa-circMLLT3 | 5.7 | 0.12 | - | - |
| **Down-regulated** |  |  |  |  |
| Hsa-circHDAC11 | 0.38 | 0.07 | - | - |
| Hsa-circFBN3 | 0.23 | 0.35 | - | - |
| Hsa-circTRPM7 | 0.41 | 0.26 | - | - |
| **Hsa-circGLCE** | **0.12** | **0.001\*\*** | **0.14** | **0.003\*\*** |
| Hsa-circFAM172A | 0.22 | 0.56 | - | - |
| Hsa-circKIF17 | 0.19 | 0.13 | - | - |
| **Hsa-circDHX8** | 0.16 | **0.008\*\*** | 0.15 | 0.29 |

IDD: intervertebral disc degeneration; Hsa: human; \*\* P < 0.01.

**Fig. 1.**
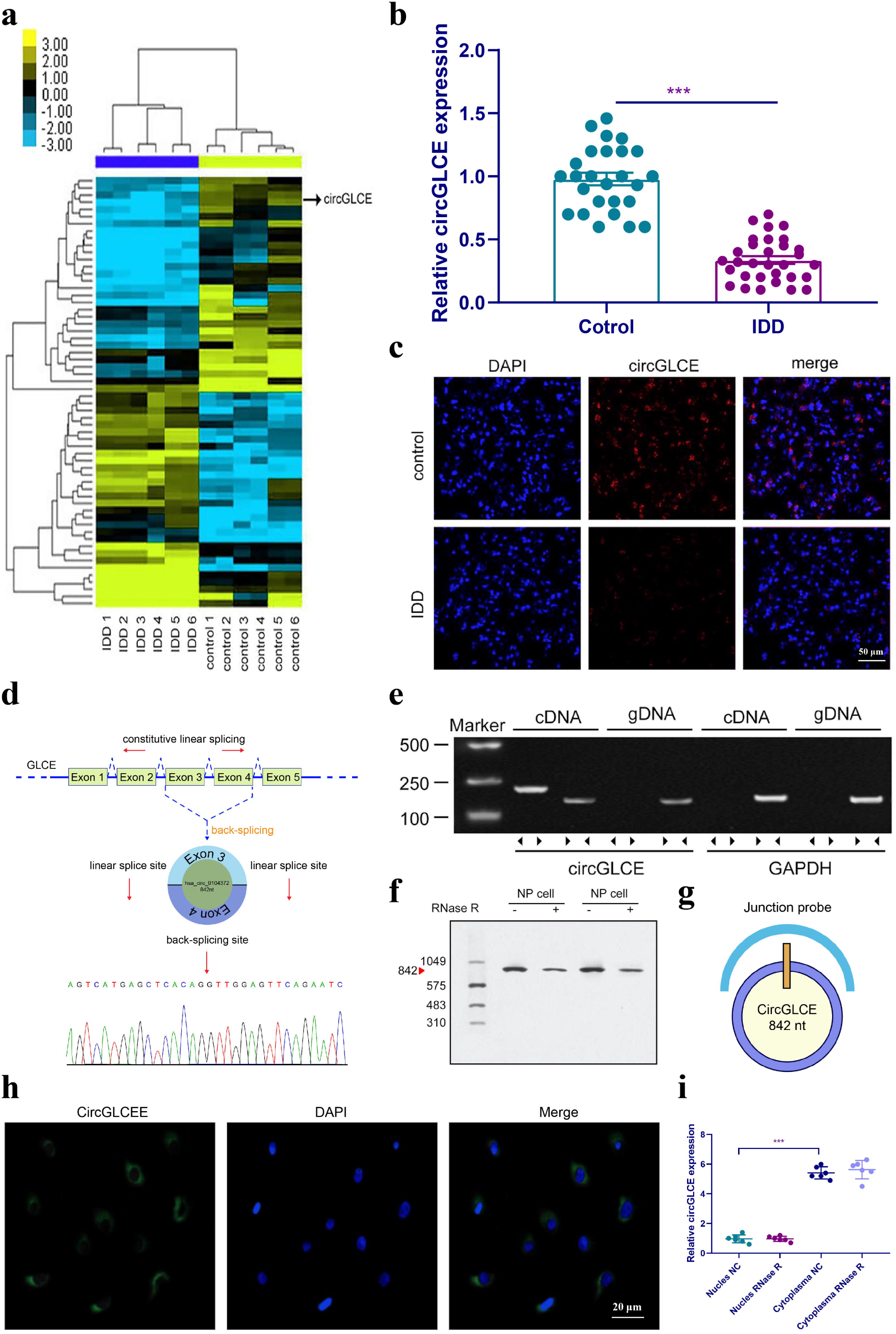
CircGLCE expression in IDD and normal NP tissue samples. (a) Heat map of all differentially expressed circRNAs between the IDD sample and the paired controls. (b) The comparison of CircGLCE expression between 31 degenerative NP samples and 26 controls, which was quantified with RT-qPCR (***P < 0.001, unpaired two-tailed Student’s t test). (c) The CircGLCE level was measured by FISH, and it also showed a significant decrease in the levels in degenerative NP samples compared to controls. (d) Schematic illustration showing the circularization of GLCE exons 3-4 from circular RNA. The Sanger sequencing of CircGLCE relied on the validation of content by RT-qPCR. (e) CircGLCE was amplified by divergent primers with cDNA. (f) CircGLCE showed RNase R resistance in northern blotting. (g) The length of CircGLCE was 842 nt. The probe targeted the junction. (h) The FISH results demonstrated the presence of CircGLCE in the cytoplasm of NP cells. (i) The results of RT-qPCR confirmed the presence of CircGLCE in the cytoplasm of NP cells (n=6, *** P<0.001, unpaired two-tailed Student’s t test). Data are the mean□±□SEM.

### 3.2 Knockdown of CircGLCE promotes NP cell apoptosis and induces the expression of matrix-degrading enzymes in NP cells

To explore the regulatory functions of CircGLCE, in vitro transfection was conducted with a CircGLCE small interfering RNA (siRNA) or a CircGLCE lentivirus for silencing or overexpressing CircGLCE, respectively. After silencing CircGLCE, which was validated with RT-qPCR (Fig. 2a), the rate of apoptosis in NP cells was remarkably increased, which was detected by flow cytometry (Fig. 2b). In comparison, overexpressing CircGLCE, which was validated with RT-qPCR (Fig. 2c), resulted in an increased rate of apoptosis in NP cells that was notably reversed by IL-1β treatment (Fig. 2d). Furthermore, according to the immunofluorescence results, by silencing CircGLCE, the expression levels of MMP13 and ADAMTS4 were increased, while COL II and aggrecan were downregulated (Fig. 2e and 2f). Overall, the silencing of CircGLCE resulted in increased apoptosis and matrix degradation in NP cells, which suggested a regulatory function for CircGLCE on those biological processes.

**Fig. 2.**
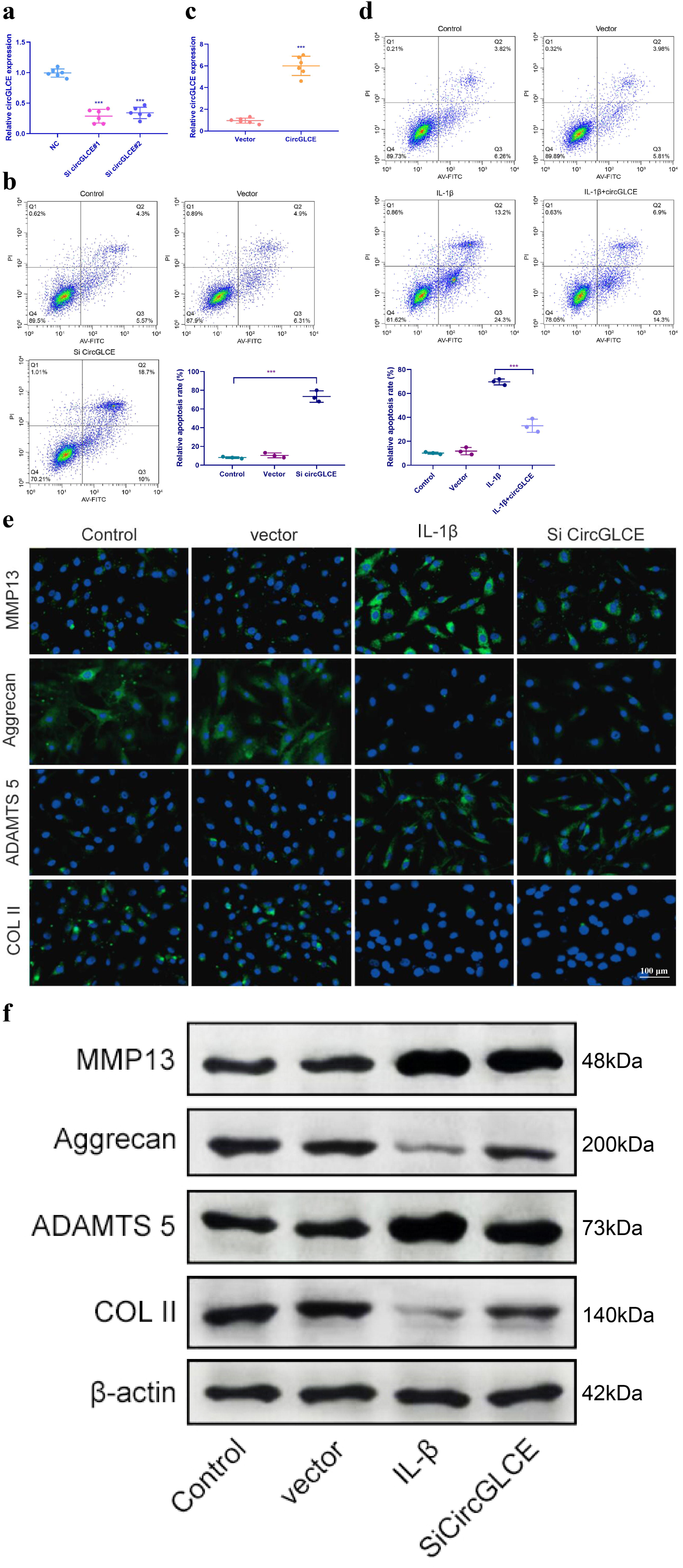
Knockdown of CircGLCE induces apoptosis and the upregulation of matrix-degrading enzymes in NP cells. (a) Forty-eight hours after transfecting NP cells with CircGLCE siRNA or a negative control, RT-qPCR showed that CircGLCE expression was effectively downregulated by silencing (n=6, ***P < 0.001, unpaired two-tailed Student’s t test). (b) Flow cytometry showed an increased ratee of apoptosis in NP cells in response to silencing CircGLCE, which was indicated by increased accumulation of signals in Q2 (n=3, *** P<0.001, unpaired two-tailed Student’s t test). (c) NP cells were transfected with CircGLCE lentivirus or the corresponding negative control, and RT-qPCR showed that CircGLCE expression was effectively upregulated (n=6, ***P < 0.001, unpaired two-tailed Student’s t test). (d) Flow cytometry showed that overexpressed CircGLCE partially reversed the apoptosis of NP cells that was promoted by treatment with IL-1β (n=3, *** P<0.001, unpaired two-tailed Student’s t test). (e) Shown by immunofluorescence analyses, silencing CircGLCE resulted in increased expression of MMP13 and ADAMTS as well as decreased expression of COL II and Aggrecan. (f) Western blot analysis confirmed the results in (e). Data are the mean□±□SEM.

### 3.3 CircGLCE serves as a miR-587 sponge in NP cells

The miRNA targets of CircGLCE were next sought out in this study due to the stable existence of CircGLCE in the cytoplasm of NP cells. First, relevant miRNAs were identified by high-throughput sequencing of the differentially expressed miRNAs between the IDD samples and normal controls, as shown in Fig. 3a. Second, relying on the analytical function of miRanda and TargetScan, miRNA recognition elements (MREs) were predicted. By comparing the MREs with the sequences of miRNAs, 7 candidate miRNAs were selected for analysis with an RNA pull down assay, and miR-587 was effectively pulled down by the CircGLCE probe (Fig. 3b). In addition, RT-qPCR results confirmed the upregulation of miR-587 in IDD samples (Fig. 3c), which was consistent with the altered CircGLCE expression due to IDD. The binding site detected by the bioinformatic approach indicated direct binding between CircGLCE and miR-587 (Fig. 3d). As shown in Fig. 3e, the dual luciferase assay demonstrated the direct binding between wild-type CircGLCE and miR-587. Accordingly, miR-587 was selected for further analysis. As shown in Fig. 3f, the colocalization of CircGLCE and miR-587 was visualized with FISH. Moreover, the regulation of ECM proteins by miR-587 was also consistent with those of the corresponding CircGLCE expression pattern. In other words, as shown in Fig. 3g and 3h, the overexpression of miR-587 enhanced MMP13 and ADAMTS5 expression but inhibited COL II and aggrecan expression, which was consistent with the effects of downregulated CircGLCE expression in IDD. In contrast, the inhibition of miR-587 caused the opposite changes in the levels of those enzymes.

**Fig. 3.**
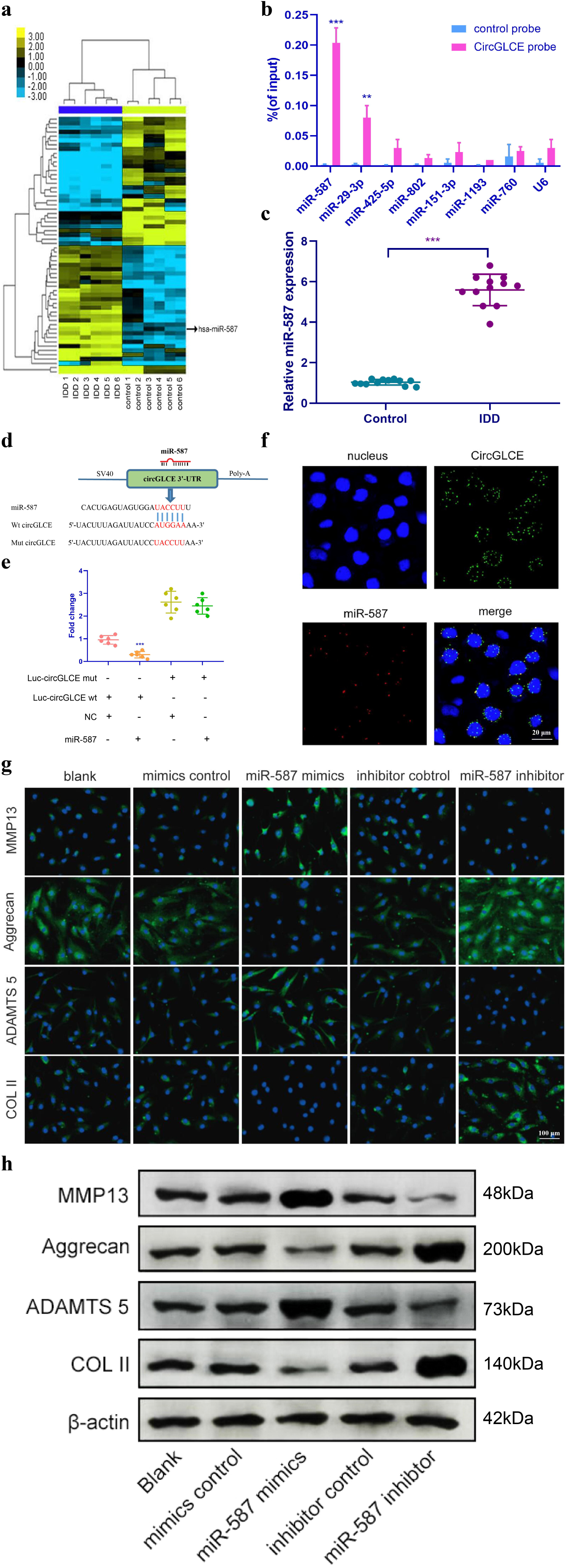
CircGLCE serves as a miR-587 sponge in NP cells. (a) Differentially expressed miRNAs in degenerative NP samples were identified with high-throughput sequencing. (b) Following RNA pull-down experiments performed with a CircGLCE probe, RT-qPCR indicated remarkably high miR-587 levels. The relative levels of CircGLCE were normalized to the input levels (***P < 0.001, one-way ANOVA coupled with Tukey’s post hoc test). (c) RT-qPCR demonstrated that miR-587 was upregulated in IDD (n=12, **P < 0.01, and ***P < 0.001; unpaired two-tailed Student’s t test). (d) The binding sites in CircGLCE for miR-587 were analysed by a bioinformatic approach. (e) NP cells were cotransfected with miR-587 mimics/negative controls and luciferase reporter constructs for wild-type or mutant CircGLCE. Dual luciferase assays demonstrated binding between miR-587 and CircGLCE (n=6, *** P<0.001, unpaired two-tailed Student’s t test). (f) FISH showed the colocalization of CircGLCE and miR-587 in NP cells. The miR-587 probe was labelled with Alexa Fluor 488, the CircGLCE probe was tagged with Cy3, and the nuclei were stained with DAPI. (g) The ECM enzymes were effectively regulated by overexpressing or knocking down miR-587. *** P<0.001. (h) Western blot analysis confirmed the results in (g). Data are the mean□±□SEM.

### 3.4 Inhibiting miR-587 counteracts the IDD-enhancing effect caused by silencing CircGLCE

To clarify the effect of the interaction between CircGLCE and miR-587 in IDD, cotransfection with sh-CircGLCE and a miR-587 sponge adenovirus was conducted, as well as transfection with wither sh-CircGLCE or the control. According to the results of western blotting (Fig. 4a), FISH (Fig. 4b) and RT-qPCR experiments (Fig. 4c), the NP cells transfected with sh-CircGLCE alone had upregulated MMP13 and ADAMTS5 expression, while COL II and aggrecan were downregulated. Nonetheless, the NP cells cotransfected with sh-CircGLCE and the miR-587 sponge adenovirus showed a reversal of the abovementioned changes because the miR-587 sponge served as a competitive inhibitor of miR-587. These in vitro analyses confirmed that the interaction of CircGLCE and miR-587 had certain regulatory effects on IDD-related matrix degradation processes.

**Fig. 4.**
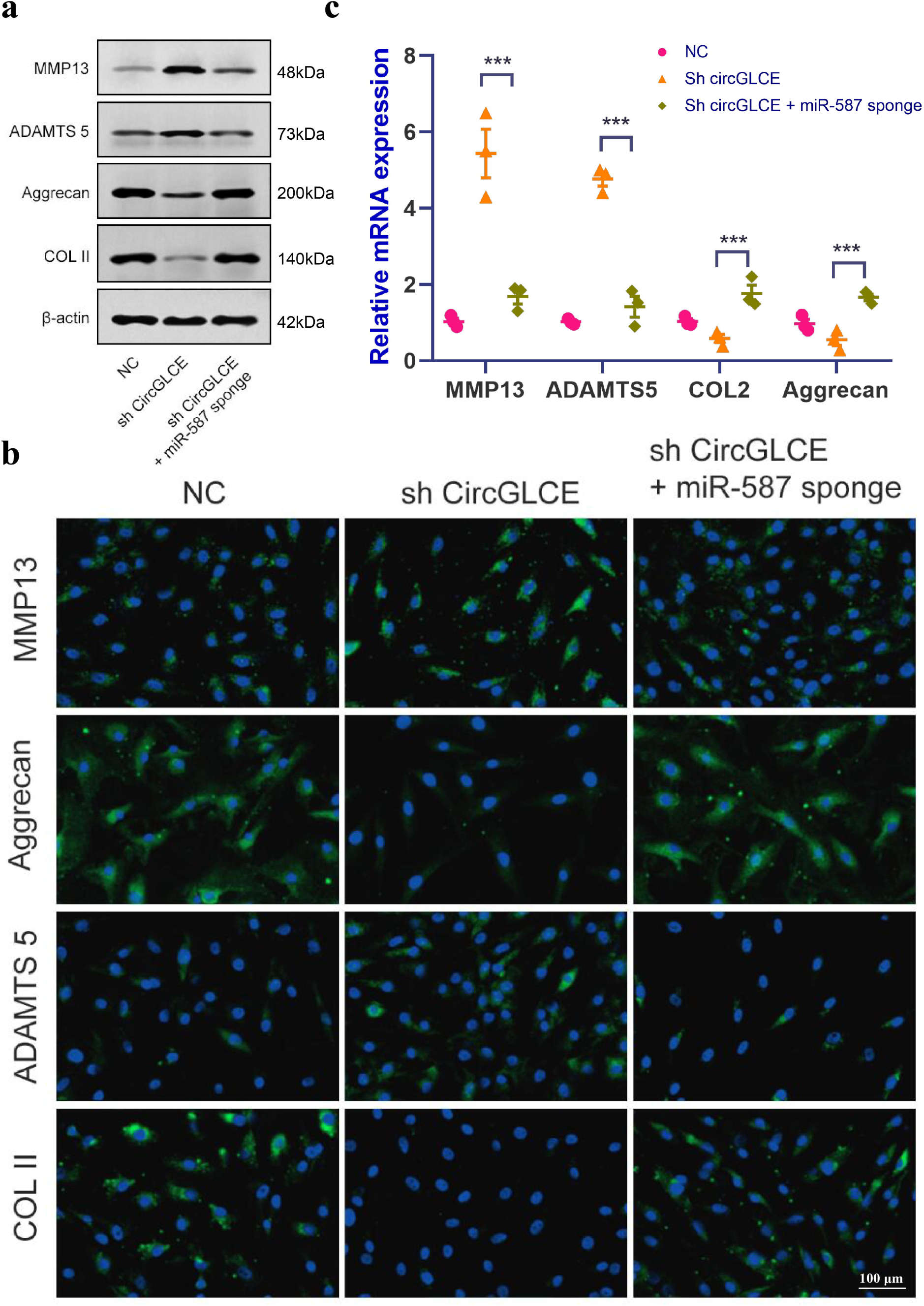
Silencing miR-587 reverses disc degeneration induced by CircGLCE downregulation. After transfecting NP cells with sh-CircGLCE, the combined sh-CircGLCE and miR-587 sponge adenovirus, or the control, (a) the levels of ECM markers were measured with western blotting; (b) the levels of ECM markers were measured with FISH; (c) the expression levels of ECM markers were measured with RT-qPCR (n=3, ***P < 0.001, one-way ANOVA coupled with Tukey’s post hoc test). Data are the mean□±□SEM.

### 3.5 STAP1 is targeted by miR-587 and mediates the function of miR-587 in NP cells

Based on prediction generated by TargetScan, miRDB and RNA-seq, the expression profiles of selected miRNAs were compared between CircGLCE-silenced samples and the paired controls by microarray analysis, as shown in Fig. 5a. Consequently, 8 candidate mRNAs were selected, among which STAP1 showed remarkably different expression patterns; therefore, we selected it for deeper research in this study. By RT-qPCR, the expression of STAP1 was found to be remarkably decreased due to degenerative disc disease (Fig. 5b), thereby suggesting that STAP1 could be inhibited by silencing CircGLCE or by overexpressing miR-587. The prediction of miRNA targets made by bioinformatic software also supports the direct binding of CircGLCE with miR-587 (Fig. 5c). Therefore, this hypothetical binding was checked with a dual luciferase assay, and the results also conformed with the interaction between miR-587 and STAP1 (Fig. 5d). Furthermore, it was confirmed that overexpressed miR-587 caused decreased STAP1 expression at both the mRNA and protein levels; in contrast, silenced miR-587 caused increased STAP1 expression (Fig. 5e and 5f). Furthermore, to determine the effects STAP1 on miR-587 in regards to the expression of IDD-related enzymes, a rescue experiment was conducted by overexpressing miR-587 alone and overexpressing both miR-587 and STAP1. The overexpression of STAP1 effectively counteracted the effects of overexpressed miR-587 (Fig. 5g and 5h), which were confirmed by western blot (Fig. 5j). As shown in Fig. 5i, STAP1 downregulated the expression of ADAMTS-5 and MMP13 and upregulated the expression of Collagen II and Aggrecan. Overall, STAP1 mediated the regulatory functions of miR-587 in IDD.

**Fig. 5.**
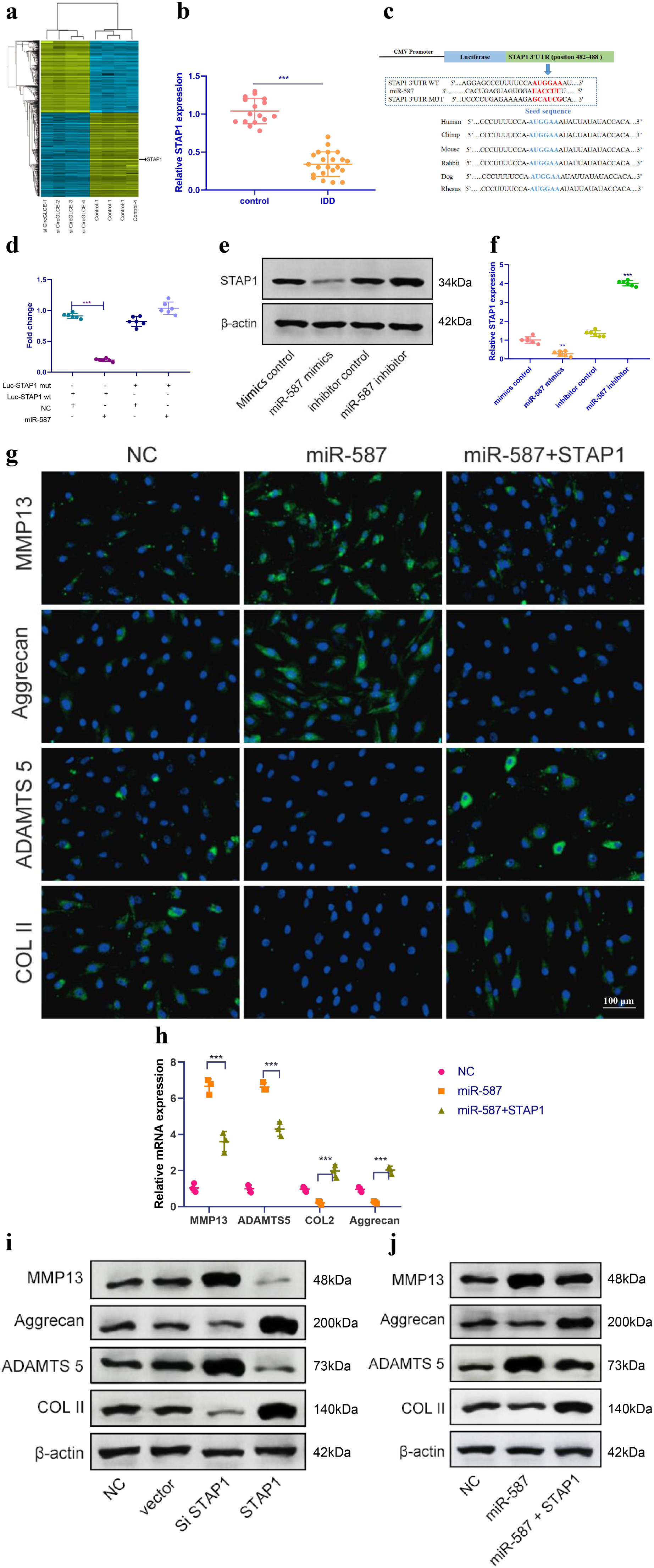
STAP1 is directly targeted by miR-587 and serves as a regulator in IDD. (a) Heat map of all differentially expressed mRNAs between the sample with CircGLCE knockdown and the paired controls. (b) The RT-qPCR results showed that STAP1 was downregulated in degenerative NP tissue samples (degenerative vs. normal=16: 22, ***P < 0.001, unpaired two-tailed Student’s t test. (c) Schematic representation of STAP1 3’-UTR demonstrating putative miRNA target site, luciferase activities of wild-type (WT-UTR), and mutant (MUT-UTR) constructs. (d) After cotransfection with the miR-587 mimics or control and the luciferase reporter constructs for wild-type or mutant 3′-UTR of STAP1, the binding of STAP1 and miR-587 was detected by dual luciferase assays (n=6, *** P<0.001, unpaired two-tailed Student’s t test. (e) The altered expression patterns of STAP1 protein were caused by miR-587 overexpression or silencing, which was detected by western blotting. (f) The altered expression patterns of STAP1 mRNA were caused by miR-587 overexpression or silencing, which was detected by RT-qPCR (n=6, **P < 0.01, and ***P < 0.001; unpaired two-tailed Student’s t test. (g) The FISH results of the rescue experiment using the pre-miR-587 adenovirus, STAP1 and the control. (h) The results of RT-qPCR confirmed the FISH results in (g). (n=3, ***P < 0.001, one-way ANOVA coupled with Tukey’s post hoc test). (i) Western blot analysis suggested that STAP1 downregulated the expression of ADAMTS-5 and MMP13 and upregulated the expression of Collagen II and Aggrecan. (j) Western blot analysis confirmed the results in (g and h). Data are the mean□±□SEM.

### 3.6 Overexpressed CircGLCE alleviates IDD in vivo

Depending on the regulatory effect of CircGLCE on IDD-related processes in vitro, it was suggested that CircGLCE could also effectively alleviate the progression of IDD in vivo. To test the function of CircGLCE, an IDD rat model was established. For adenoviral CircGLCE treatment, the wild type was expected to induce the same biological effect as endogenous CircGLCE overexpression, while the mutant type was expected to have no biological effect. Twelve weeks after the administration of wild-type or mutant CircGLCE to the IDD groups, the group with overexpressed CircGLCE showed a significantly increased disc height index (%) compared to the untreated and mutant-CircGLCE-treated IDD+ groups (Fig. 6a and 6b). Additionally, the Pfirrmann grading of IDD in MRIs showed a consistent pattern, which further confirmed that the degree of IDD was reduced by overexpressed CircGLCE in vivo (Fig. 6c). Moreover, at the cellular level, the overexpression of CircGLCE resulted in miR-587 expression being remarkably reduced in the NP sample of the IDD model, as shown in Fig. 6d and 6e. Accordingly, overexpressed CircGLCE inhibited IDD-related cellular activities, such as apoptosis of NP cells, as indicated by TUNEL staining and RNA FISH (Fig. 6f and 6g). Additionally, ECM degradation was effectively prevented by CircGLCE expression, as shown by western blotting, H&E staining, and immunohistochemical assays (Fig. 6h and 6i). The histological findings in the IDD group were a loss of cells in the NP region of intervertebral disc. Increased numbers of NP cells were observed in the group treated with circGLCE compared to the IDD group. The expression of MMP13 was higher in the IDD group than it was in the IDD++circGLCE group. In conclusion, overexpressed CircGLCE alleviated IDD in vivo by suppressing catabolic cellular activities through acting as a miR-587 sponge. The results of histological examination were consistent with these observations (Fig. 6j).

**Fig. 6.**
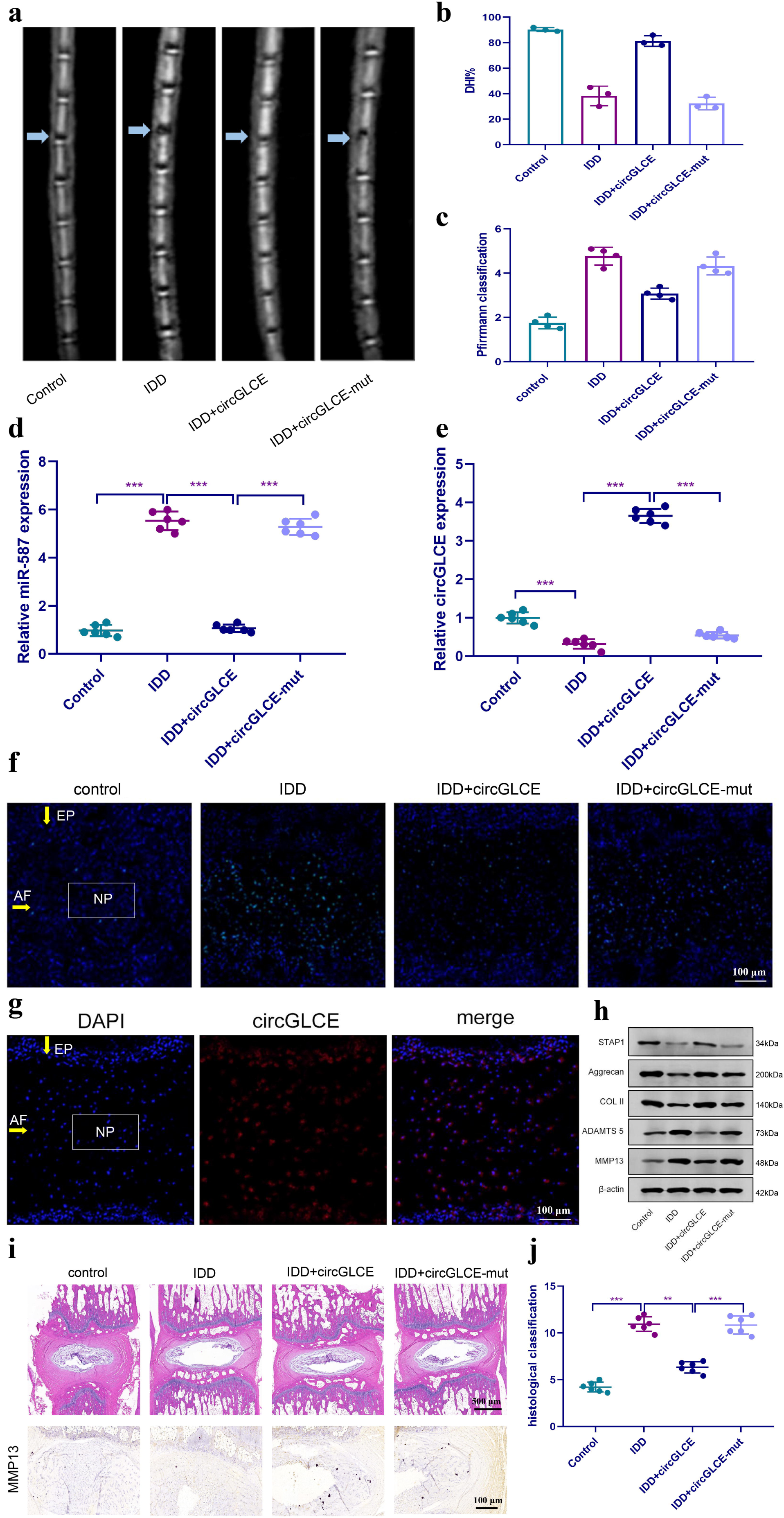
Overexpressed CircGLCE alleviates IDD in vivo. 12 weeks after the induction of IDD and the administration of treatments, (a) MRI was performed and images were obtained; (b) the disc height indices after (n=3,*** P<0.001, unpaired two-tailed Student’s t test); (c) the grading of IDD based on Pfirrmann classification (n=3, *** P<0.001, unpaired two-tailed Student’s t test); (d) the expression of miR-587 was downregulated by overexpressing CircGLCE in an animal model of induced IDD, as shown with RT-qPCR (n=6, ***P < 0.001, unpaired two-tailed Student’s t test); (e) RT-qPCR results indicated the CircGLCE levels corresponding to induced IDD and CircGLCE treatments (n=6, ***P < 0.001, unpaired two-tailed Student’s t test); (f) TUNEL staining revealed the in vivo effect of overexpressed CircGLCE on apoptosis after 12 weeks; (g) RNA FISH illustrated the in vivo distribution of CircGLCE in NP tissue; (h) western blotting showed the expression profiles of ECM markers corresponding to induced IDD and CircGLCE treatments; (i) H&E and immunohistochemistry staining of NP tissues. The cells of the NP region in the intervertebral disc were more abundant in the control and IDD+circGLCE groups than they were in the IDD and IDD+circGLCE-mut groups. The expression of MMP13 was higher in the IDD group than it was in the IDD++circGLCE group. (j) Histological examination (n=6, ** P<0.01, and *** P<0.001; unpaired two-tailed Student’s t test). Data are the mean□±□SEM.

## 4. Discussion

NP cells are critical for maintaining the structural integrity of intervertebral discs,^5, 6^ so the dysregulation of cellular activities closely correlates with the pathogenetic mechanism of IDD;^7^ however, intensive research work is still required to facilitate the improvement of prophylactic and curative approaches.^38^ Recently, it has been reported that specific circRNAs have been identified as significant regulators and biomarkers for degenerative disc disease.^39-42^ The underlying mechanism mostly relies on their abilities to counteract the inhibitory effects of miRNAs on the target genes. Due to the development of microarray profiling, bioinformatic approaches and sequencing techniques, the identification of novel circRNAs as effective regulatory factors and potential therapeutic targets for IDD is expected.^43^

CircGLCE is highly abundant in the mammalian brain and is especially enriched in the cerebellum.^44^ Due to its high stability, it might be used by neuronal termini and molecular postsynaptic platforms as a synaptic tag to maintain molecular memory. By detecting the differentiated circRNA expression profiles between degenerative NP samples and paired normal tissues with microarray, CircGLCE was selected for further exploration in this study due to its significance in IDD progression. In vitro RT-qPCR results confirmed that CircGLCE was downregulated in IDD samples. Moreover, the FISH results demonstrated the presence of CircGLCE in the cytoplasm of NP cells. To clarify the effect of decreased CircGLCE expression, additional in vitro assays of IDD-related activity were conducted upon the knockdown of CircGLCE in normal NP tissues. The results suggested that decreased CircGLCE expression was associated with increased apoptosis of NP cells and enhanced ECM degradation, which are key biological processes contributing to the development of IDD. To further validate the function of overexpressed CircGLCE, IL-1β treatment was applied to induce a significant increase in apoptosis,^45-47^ and this effect was partially reversed by CircGLCE overexpression. Enhanced ECM degradation was demonstrated by decreases in ECM components, namely, Collagen II and aggrecan, as well as by increases in ECM-related proteinases, namely, MMP-13 and ADAMTS5.^48, 49^ Therefore, the involvement of CircGLCE in the pathogenesis of IDD was proven.

Depending on the sequence, splicing and circularization of CircGLCE determined in this study, the predominant direct target was predicted to be miR-587, which was validated by the RNA pull-down assay. According to the main regulatory mechanism of circRNAs, in which they serve as miRNA sponges,^17, 20^ CircGLCE was predicted to suppress the function of miR-587 by binding to it. The results of this study showed that the increased miR-587 level detected by RT-qPCR in IDD samples was consistent with the decreased CircGLCE in IDD. Moreover, the binding and colocalization of CircGLCE and miR-587 was confirmed. To clarify the effect of miR-587 on ECM degradation, by either silencing or overexpressing miR-587, the expression levels of ECM-degrading markers effectively responded to altered miR-587 levels. Accordingly, the results of in vitro experiments indicated that the inhibition of miR-587 exerted by CircGLCE prevented ECM degradation. Although another study reported that miR-587 had a certain effect on apoptosis-related mechanisms,^50^ this was the first study to point out its role in matrix degradation.

With a similar bioinformatic approach, the target of miR-587 was identified. The decreased STAP1 expression in IDD was consistent with the downregulation of CircGLCE and the increase in miR-587 activities, which resulted in enhanced matrix degradation. In contrast, the overexpression of STAP1 reversed the effect of increased miR-587 expression, thereby inhibiting the progression of IDD. This was also the first study to identify STAP1 as part of the regulatory mechanism for IDD, while it has been generally recognized as a critical factor in lipid catabolism.^51^ Furthermore, it was suggested that STAP1 could directly regulate the expression of Collagen II, Aggrecan, ADAMTS-5 and MMP13. However, the study on the function of STAP1 in IDD is part of a separate research project, and the data are not yet published.

The clinical significance of CircGLCE and its downstream pathways could be further verified with in vivo examinations using an animal model of induced IDD. As expected, the treatment causing increased CircGLCE expression in vivo effectively alleviates the severity of IDD in terms of clinical manifestations and cellular activities. Therefore, CircGLCE is proposed as a promising therapeutic target. As effective regulatory factors in IDD, miR-587 and STAP1 might also have therapeutic potential. Nonetheless, the enclosed circular structure contributed to the stability of CircGLCE in the cytoplasm of NP cells, suggesting that targeting CircGLCE might be more efficient as a therapeutic solution.^52^ In addition, a comprehensive understanding of the relevant molecular and biological mechanisms might also contribute to treatment efficacy, such as improving the precision of targeting by therapeutic substances.^53^

In addition to the significant findings of this study, limitations on the scope of research still exist. First, although the regulation of NP cell apoptosis by CircGLCE was demonstrated, the association between its regulatory functions on apoptosis and matrix degradation was not fully clarified. Second, the downstream signalling pathway for STAP1 was not studied. The initiation of IDD, especially the molecular activity inducing the downregulation of CircGLCE, might be further investigated in follow-up research works.

To summarize, CircGLCE alleviates intervertebral disc degeneration by regulating apoptosis and matrix degradation through the targeting of miR-587/STAP1. Our study highlights the potential clinical application of a CircGLCE-based therapy to treat IDD patients. It would be important in the future to further test the combination of NP cell-specific delivery of CircGLCE (or small molecule compounds that can promote endogenous expression of CircGLCE) through sponging miR-587 and thereby derepressing STAP1 to achieve an optimal anti-IDD effect with minimal side effects. The findings expand the current understanding of IDD pathogenesis while identifying novel promising therapeutic targets for the development of IDD treatments.

## Acknowledgements

This work is supported by The National Science Foundation of China (No. 81802203). The funders had no role in study design, data collection, data analysis, interpretation, or writing of this report.

## Author contributions

CZH, ZWB, and LYM designed the experiments. ZWB and DM collected samples. CZH, ZWB and DM conducted the experiments and acquired the data. CZH, ZWB, DM and ZY analysed the data. CZH wrote the manuscript. ZWB, DM, ZY and LYM revised the manuscript. All authors approved the final version. The corresponding author had full access to all the data in the study and had final responsibility for the decision to submit for publication.

